# Long-Term Dynamics of Esca Disease in Western Switzerland

**DOI:** 10.64898/2026.10.28.762386

**Authors:** Matthieu Wilhelm, Vinciane Monod, Anne-Lise Fabre, Olivier Viret, Vivian Zufferey, Katia Gindro, Pierre-Henri Dubuis

## Abstract

Agroscope conducted a long-term survey to assess the incidence of esca disease in Western Switzerland between 2001 and 2022 and to characterize its temporal dynamics. This study has four main objectives: (1) to provide a rare and particularly valuable long-term dataset documenting the incidence of esca over more than two decades; (2) to identify potentially important climatic drivers using a recent model-independent variable importance algorithm that quantifies the strength of their association with esca incidence; (3) to develop predictive models of esca incidence using climatic covariates derived from microclimatic weather data; and (4) to investigate the relationship between water balance and regionally aggregated esca incidence. The climatic variables identified by the variable importance algorithm as being most strongly associated with esca incidence at the plot level were primarily related to water balance.

The predictive analysis showed that model performance at the plot level was limited. Consistent with previous studies, analysis of the variables contributing to model predictions did not allow us to identify a single factor, or set of factors, that could reliably predict esca incidence.

At the regional scale, however, water balance and seasonal vapor pressure deficit (VPD) were strongly associated with esca incidence across years, with drier regions generally showing lower disease incidence.

Finally, addressing the initial question that motivated this long-term survey, we observed a slight decreasing trend in esca incidence over the study period. This trend coincided with increasingly dry conditions in Western Switzerland and may therefore be partly associated with changes in regional water balance in the context of climate change.

All the datasets and analyses presented here are made available and reproducible.

## 1 Introduction

Long-term, large-scale vineyard monitoring is essential for understanding disease dynamics, identifying spatial and temporal patterns, and supporting sustainable disease management strategies. Among the diseases affecting grapevine production worldwide, esca and other grapevine trunk diseases (GTDs) remain among the major challenges facing viticulture. These diseases cause substantial yield losses, vine decline and mortality, leading to significant economic consequences (Bertsch et al., 2013; Gramaje et al., 2018; Hofstetter et al., 2012). Across Europe, esca has been increasingly reported over recent decades. (Bruez et al., 2012; Dewasme et al., 2022; Etienne et al., 2024) A similar increase has been reported globally (Kenfaoui et al., 2022).

Esca is a complex disease syndrome characterized by necrotic lesions in woody tissues and foliar symptoms that can culminate in complete canopy loss (Fischer and Peighami-Ashnaei, 2019; Ouadi et al., 2019). Despite extensive research, the mechanisms underlying symptom expression remain incompletely understood. In particular, the precise role of pathogenic microorganisms in foliar symptom development is still debated (Bruez et al., 2020; Del Frari et al., 2019; Monod et al., 2025a). Current evidence suggests that symptom expression results from interactions among multiple biotic and abiotic factors acting as predisposing conditions or triggering events. Host-related factors reported to influence disease expression include cultivar susceptibility (Chacón-Vozmediano et al., 2021; Gastou et al., 2024; Guerin-Dubrana et al., 2019), vineyard age (Etienne et al., 2024; Fussler et al., 2008), and nitrogen fertilization practices (Dell’Acqua et al., 2025). Environmental conditions play an equally prominent role. Esca incidence has been linked to temperature regimes, precipitation patterns, drought events, and other climatic variables affecting plant water status and physiological functioning (Bortolami et al., 2021; Calzarano et al., 2018; Serra et al., 2018; Surico et al., 2000). Likewise, soil characteristics influencing water availability have been associated with increased symptom expression (Monod et al., 2025b). Collectively, these findings highlight the importance of plant-environment interactions in shaping annual esca incidence.

A major challenge in studying esca epidemiology is the strong year-to-year variability in symptom expression. Esca incidence can fluctuate substantially between consecutive seasons (Calzarano et al., 2018; Etienne et al., 2024, 2026), making short-term observations potentially misleading when assessing long-term disease dynamics. Consequently, long-term monitoring is necessary to distinguish transient fluctuations from genuine epidemiological trends. Furthermore, although climatic factors are frequently associated with symptom expression, their ability to predict disease incidence across sites and years remains insufficiently explored.

To address these questions, we conducted a 21-year survey of a network of vineyard plots representing ten grapevine cultivars across six viticultural regions of Western Switzerland . Esca foliar symptom incidence was monitored annually or biennially throughout the study. This long-term dataset enables the investigation of temporal and spatial patterns of esca expression and the assessment of the predictive value of meteorological variables at the plot level. We selected climatic indicators known to influence grapevine physiology and previously associated with esca incidence, including temperature, atmospheric pressure, relative humidity, radiation, precipitation, evapotranspiration, and vapor pressure deficit. Using a predictive modelling approach applied to climatic time-series data, we tested the extent to which these environmental variables could predict esca incidence at both plot and regional scales. We hypothesized that climatic variables affecting grapevine water balance would explain a significant proportion of interannual variation in esca incidence and could therefore help predict disease occurrence across vineyards and regions.

## 2 Materials and methods

The data collection was not initially designed as a long-term monitoring program, nor was the sampling scheme intended to provide comprehensive coverage of all viticultural regions in Western Switzerland or to follow a fully randomized, reproducible sampling design. Some of the variables collected were not used in the analyses, essentially because over the years they were found to be unreliable or irrelevant (and are thus missing for the final years): an estimated age category and the vine training system. From a statistical point of view, the dataset is therefore a convenience sample rather than a well-designed survey.

### 2.1 Dataset

Vineyards were selected within six viticultural areas: La Côte, Lavaux, Chablais, Nord-Vaudois, Valais and, from 2012 onward, Neuchâtel. These regions encompass contrasting climatic conditions, particularly with respect to rainfall patterns. In 2001, a total of 66 parcels were surveyed on La Côte, 15 in Lavaux, 4 in the Nord Vaudois and 2 in Valais. Additional vineyards were added in 2004 to reach 21 parcels in Valais, and from 2006 the sampling was homogenized across regions (31 parcels on La Côte, 12 in Lavaux, 12 in Chablais and 17 in Valais). In 2012, nine Neuchâtel vineyards were integrated into the network, along with seven additional chasselas plots in Valais to distinguish cultivar effects from local environmental factors. Plot coordinates were systematically recorded. When a vineyard was uprooted or showed excessive missing vines, it was replaced by another nearby plot of the same cultivar. Over the full monitoring period, 31 plots remained continuously in the survey (15 from 2004 at least, 16 from 2012), including 24 planted with Chasselas. The rootstock used was primarily 3309C. It should also be noted that there is no irrigation on the plots examined.

Each year, symptom incidence was carried out between late August and mid-September, when seasonal foliar symptoms are fully expressed. Experienced observers assessed four groups of 100 vines per plot (400 vines in total) for esca incidence. The following categories were recorded: foliar symptoms of slow esca, apoplectic form, eutypa dieback, and vines missing, recently replaced (1–2 years), or not budded. Disease incidence is quantified as the percentage of vine stocks showing foliar symptoms or apoplexy within each plot. We use the sum of the percentages of plants affected by symptoms and those that suffered apoplexy as the response variable for all analyses.

From 2004 onward, surveys were repeated biennially, and annually from 2014 until monitoring ended in 2022.

#### 2.1.1 Stable plots

A subset of “stable plots” — plots that were consistently monitored over multiple years — was identified to allow longitudinal analyses controlling for site-specific effects. Note that this subset cannot be inferred from the identifier of the plot alone. Indeed, the plot identifier indicates a location or a domain rather that a specific plot. Two stability groups are defined:

• **2004–2022**: plots present at every observation campaign from 2004 to 2022 (sometimes even from 2001, but we restricted to 2004 to ensure a minimum number of complete observations). There are 15 plots in this group.
• **2012–2022**: plots present from 2012 to 2022 (larger subset with annual observations). There are 16 plots in this (in addition to the previous group).

These subsets enable a within-plot temporal comparison, but the number of plots of those categories is rather limited.

### 2.2 Covariates

In addition to the response variable, a set of covariates was collected or derived for each plot-year observation. Some covariates were directly measured in the field (location, variety etc.), while others were obtained from external sources. We essentially focused on variables that could potentially influence Esca expression, namely meteorological data and soil water reserve. Many climatical variables are reported in the literature as potential drivers of Esca expression, see e.g. Etienne et al. (2026) for an account. We did include all the ones that were available, but we stress the fact that those variables are not always faithfully representative of the microclimate of the plot, essentially because of the distance between the plot and the weather station. Over the years, the number of weather stations has increased, and the distance between the plots and their nearest station has decreased, due to the expansion of the Agrometeo network.

#### 2.2.1 Meteorological data

Climate data was obtained from two complementary networks:

• **MeteoSwiss** (Federal Office of Meteorology and Climatology): 8 stations relevant for the study area, providing complete temporal coverage from 2001 to 2022. Measured variables include temperature, atmospheric pressure, precipitation, global radiation, relative humidity, and evapotranspiration.
• **Agrometeo** (Agroscope agrometeorological network): approximately 40 stations, generally located closer to vineyard plots. The deployment of the network started in 2003 and additional stations have been added over the time. The same variables are available as MeteoSwiss, excluding atmospheric pressure and global radiation (for some of the stations).

##### Merging strategy

A “Agrometeo-first with MeteoSwiss fallback” approach was adopted. For each plot-year combination, Agrometeo data is used if the nearest station is within 30 km and data is available for that year; otherwise, MeteoSwiss data from the closest station is used. Note that we have not considered MeteoSwiss stations that are higher than the highest Agrometeo station even if they were closer, as they are not representative of the vineyard climate.

##### Temporal aggregation

All climate variables are aggregated monthly over the growing season (March–August; since the observations have been made in late August in the vast majority of cases, it is irrelevant to include climatic variables later), yielding 6 monthly values per variable type (approximately 36–56 climate predictors in total depending on available variables). Additionally, seasonal aggregates (denoted by SeasXX) summarize the entire March–August period into a single value per variable.

##### Vapour Pressure Deficit (VPD)

It is computed from hourly temperature (*T*) and relative humidity (RH, in %) data using the Magnus formula (Alduchov and Eskridge, 1996):

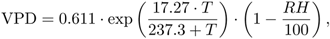

then aggregated to daily values as the maximum daily VPD and finally aggregated to monthly values as the mean of the maximum daily VPD over each month.

#### 2.2.2 Other covariates

The non-meteorological covariates that are included in the dataset are summarized in Table 1.

**Table 1.** Some of the collected covariates.

| Variable | Type | Description |
| --- | --- | --- |
| variety | Categorical | Grape variety (chasselas, gamay, pinot noir, pinot gris, sauvignon blanc, and minor varieties). |
| RU_nb | Numeric | Available water reserve from soil surveys (mm). Only available for 25 stable plots. |

Regarding the covariate RU_nb (available water reserve), it is only available for a limited number of plots. The RU is an estimate based on the soil survey and should therefore be interpreted with caution. The available water reserve has been determined based on a previous study of the wine-growing land (“terroirs”) of Canton de Vaud (Letessier and Fermond, 2004).

### 2.3 FORD algorithm

FORD (*Feature Ordering by integrated R*^2^ *Dependence*, Azadkia and Roudaki, 2025) is a variable selection algorithm. It is based on a measure of the strength of relationship between a set of predictors and the response variable using the so-called integrated *R*^2^. This dependence measure does not rely on any specific model or distributional assumptions. It can capture non-linear and non-monotonic relationships, which is in strong contrast with traditional correlation measures. The FORD algorithm adds one variable at a time to the “acting set’ ’ (the set of currently selected variables), and at each step determines which variable has the strongest measure of association with the response variable. The procedure stops when no remaining variable shows an increase in the association between the active set and the response.

### 2.4 Machine learning predictive study

Since the underlying causal mechanism of the disease remains poorly understood, it seemed more reasonable to us to use models that were as flexible as possible, at the potential cost of some interpretability. Thus, we chose a predictive approach rather than an inferential one (Breiman, 2001b). Another reason to choose a predictive approach relates to the sampling design, which was not designed to guide a specific model, as might be the case in controlled trials see e.g. (Meier, 2022).

#### 2.4.1 Data splitting, baseline, metrics and hyperparameter selection

Using a conventional practice in machine learning (Hastie et al., 2009), the dataset was split into 75% training / 25% test set and stratified by the response variable (total incidence). The stratification ensures that the distribution of the response variable is comparable in both sets. 10-fold cross-validation (single repeat) was applied on the training set for hyperparameter tuning when needed.

The means of the incidence by region and year was used as the baseline prediction for all test observations. This represents the simplest possible model, one that predicts the same value regardless of covariates (except for region and year). We used Mean Absolute Error (MAE), i.e. the average difference in absolute value, as the primary metric and the relative change of the MAE with respect to the baseline (in percent; negative values indicate an improvement over the baseline) is defined as

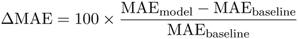

#### 2.4.2 Algorithms and methods

The pipeline was run across multiple data subsets to assess its sensitivity to different settings and plot conditions. The tested configurations are reported in Table 2.

**Table 2.**
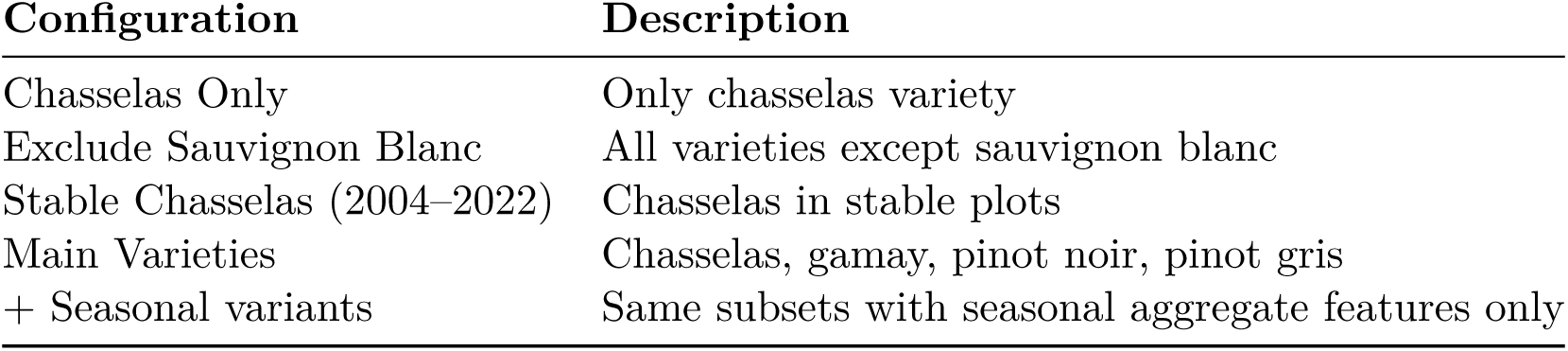
Definition of the different configurations used by the predictive models.

| Configuration | Description |
| --- | --- |
| Chasselas Only | Only chasselas variety |
| Exclude Sauvignon Blanc | All varieties except sauvignon blanc |
| Stable Chasselas (2004–2022) | Chasselas in stable plots |
| Main Varieties | Chasselas, gamay, pinot noir, pinot gris |
| + Seasonal variants | Same subsets with seasonal aggregate features only |

Each configuration was run with both monthly and seasonal aggregation modes (8 configurations total).

On each configuration, three machine learning algorithms were trained and compared:

• **Random Forest** (Breiman, 2001a); ranger engine (Wright and Ziegler, 2017)
• **Extreme Gradient Boosting** (Chen and Guestrin, 2016); xgboost engine (Chen et al., 2026
• **LASSO regression** (Tibshirani, 1996); glmnet engine (Friedman et al., 2010

Each of those algorithms comes with their own variable importance metric, which quantifies the contribution to each of the variables to the estimate. To obtain a unified ranking across models and configuration using different importance scales, the following aggregation was applied:

1. **Rescale** each model’s importance to [0, 1] by dividing by the within-model per configuration maximum
2. **Weight** each model proportionally to its MAE improvement over baseline:

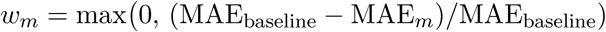

3. **Combine** via weighted average: 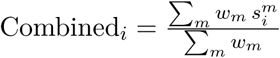

Thus, the weight assigned to a model for a given configuration in the calculation of overall importance is proportional to its predictive power (relative to the baseline). Models that do not beat the baseline receive zero weight.

### 2.5 Regional aggregation, differences and hydrology

We computed two climate indicators for each of the six study regions:

• **P/ET ratio** (precipitation/evapotranspiration) ratio — the mean annual ratio of precipitation (P) to FAO reference evapotranspiration (ET), computed from MeteoSwiss data (when available) at one representative station per region. The ratio is computed over the 12 months preceding the observations (September of previous year to August) and is a unitless measure of the water balance
• **Seasonal VPD** — the median seasonal VPD (mean of maximum daily VPD, March– August) across all plot-year observations within each region.

The P/ET ratio is scale-invariant and thus allows comparison of regions with different rainfall patterns. It is a sensible indicator of the hydrometry of a given region. The seasonal VPD was already chosen as a proxy for the water stress experienced by the plant. Both indicators were compared to the median Esca incidence (Symptoms + Apoplexy, %) by region and year. Note that these two indicators are highly correlated and thus essentially convey similar information. To compute the P/ET ratio, we used the following weather stations of the MeteoSwiss network: Aigle (Chablais), Changins (La Côte), Pully (Lavaux), Neuchâtel (Neuchâtel), Mathod (Nord Vaudois) and Sion (Valais). Note that Mathod has a few missing data.

### 2.6 Codes, data availability and reproducibility

All the codes to run the analyzes are made available on github, while the codes that were used to download, organize and preprocess the meteodata are only available on demand, since for some of them, it requires access to the API of Agrometeo. The maps are also only available on demand because they rely on external shapefiles. The data set itself is published on Zenodo (Fabre et al., 2026).

## 3 Results

### 3.1 Exploration of the dataset

The full dataset comprises 1328 plot-year observations across the 6 viticultural regions of western Switzerland as shown in the map depicted in Figure 1.

**Figure 1.**
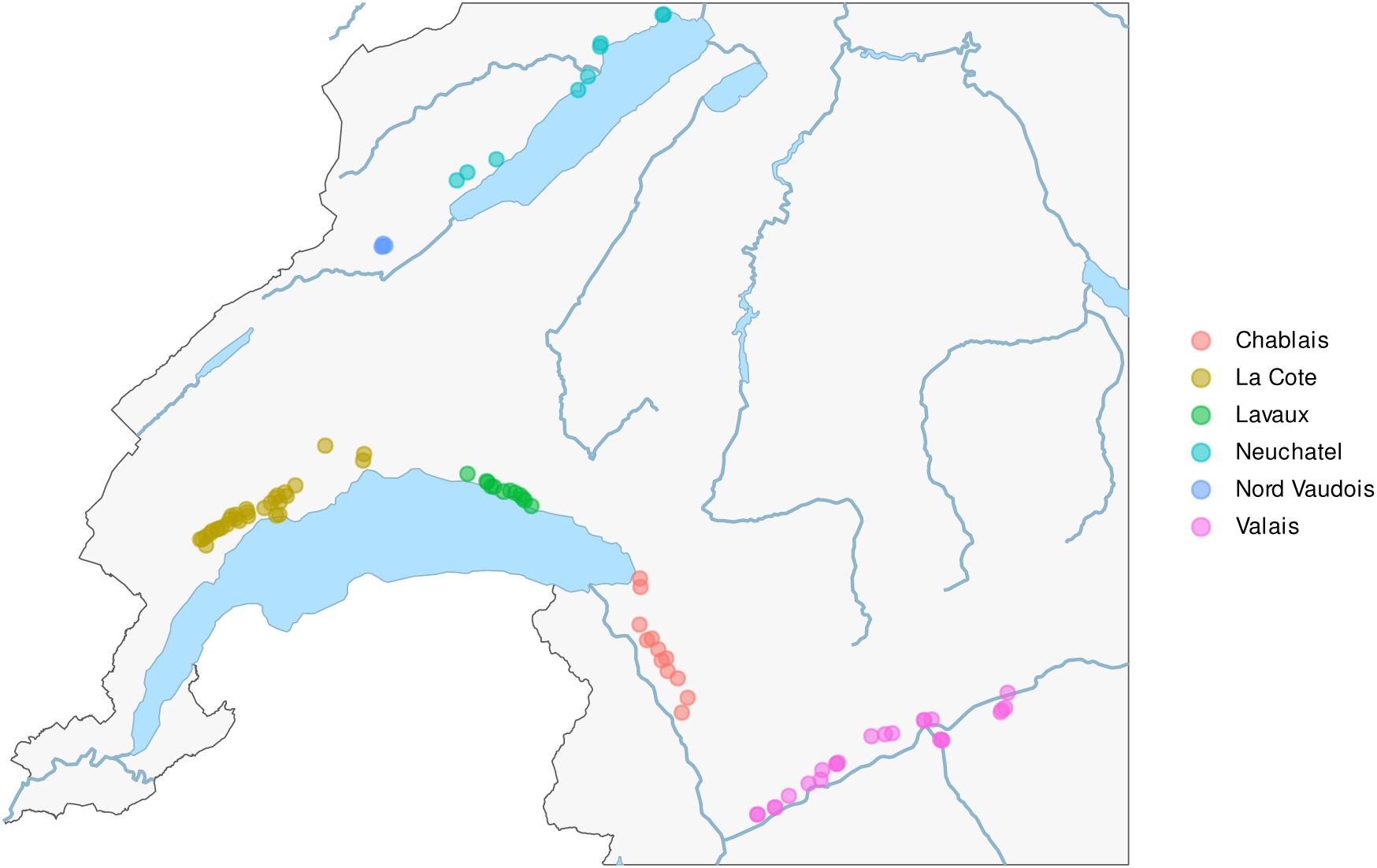
Map of the plots used in this study. An interactive version with the locations of the meteorological stations is available in the supplementary material.

The sample composition is heavily dominated by the La Côte region. On Figure 2b, the distribution of the samples in terms of cultivar and regions over the years is displayed. Chasselas is predominant in the cultivars, essentially because of its importance in Western Switzerland, while the La Côte region was the most sampled, because of proximity with the Changins agronomical research station.

**Figure 2.**
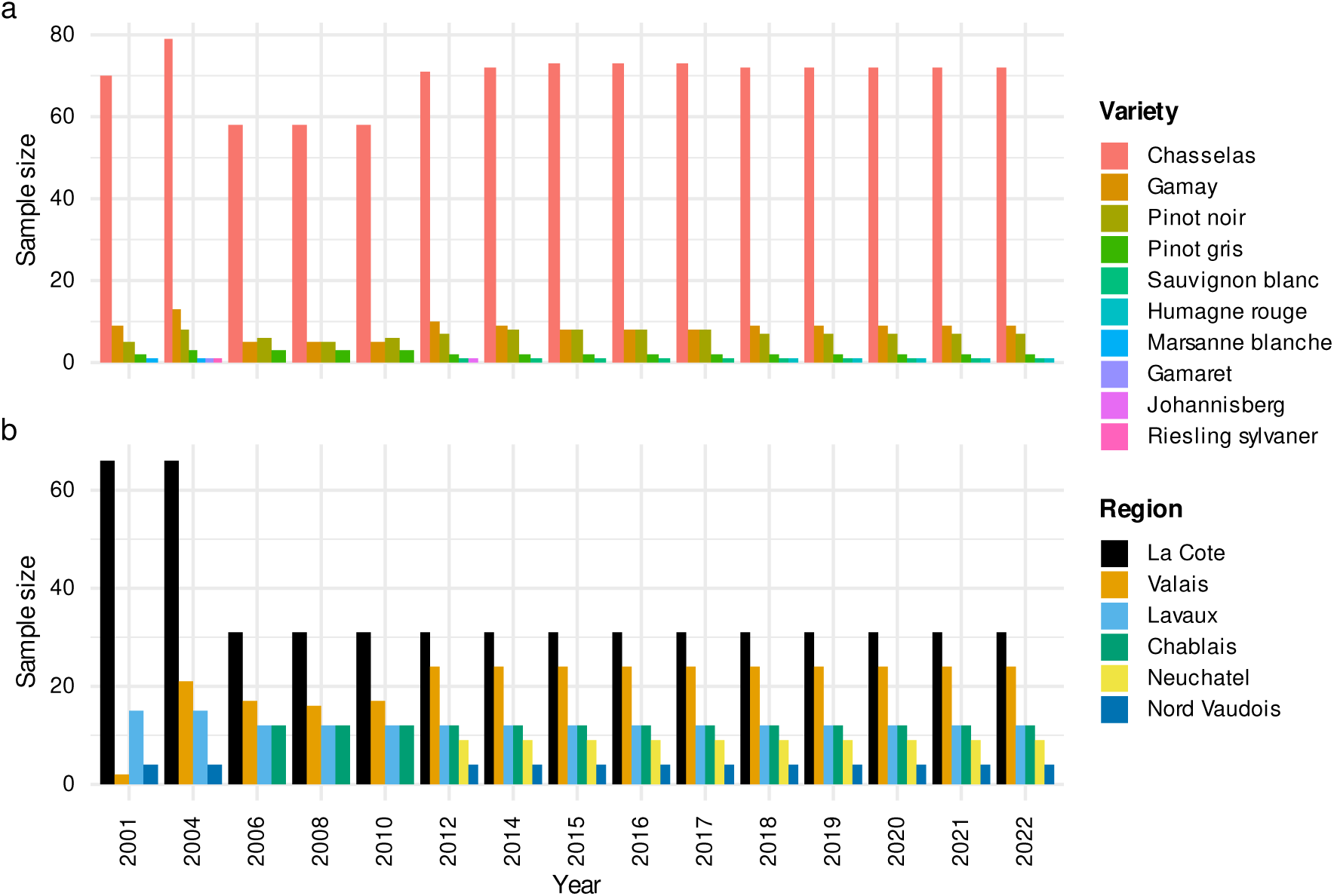
Number of plots sampled per year (a) by grape variety and (b) by viticultural region. temporal patterns. We observed a slight negative trend in the incidence of the esca disease over the years.

### 3.2 Temporal evolution of Esca expression

The temporal evolution of the combined Esca expression (foliar symptoms + apoplexy, in % of vine stocks) is displayed as yearly boxplots across all observed plots (Figure 3a). Smooth trend lines for the mean (loess) and median (Fasiolo et al., 2021a,b) are also displayed to highlight the

**Figure 3.**
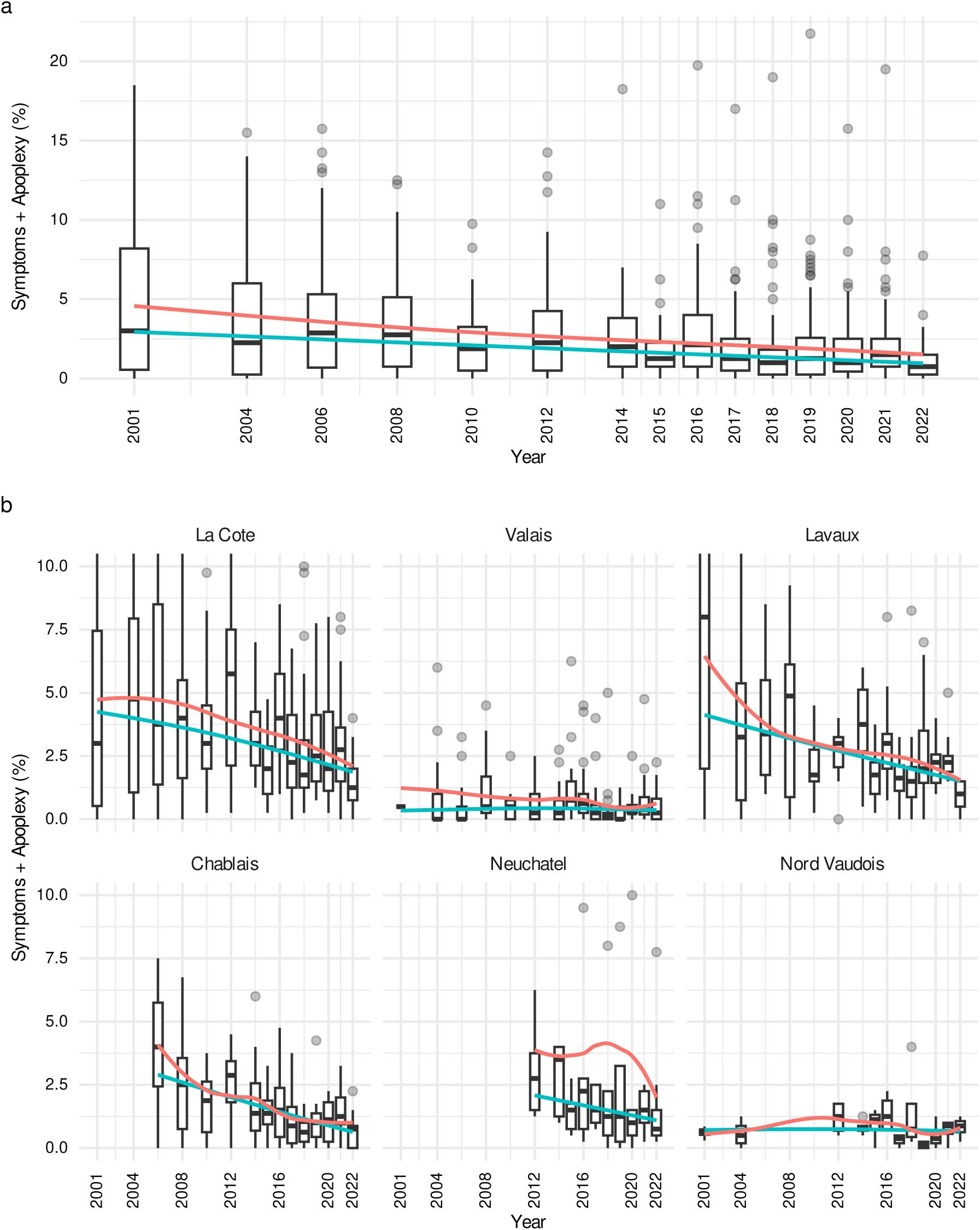
Temporal evolution of esca incidence, expressed as the percentage of vine stocks with foliar symptoms or apoplexy per plot, (a) for all plots and (b) by viticultural region. Boxplots show the distribution of plot incidences in each survey year. Red lines: smooth mean (loess); blue lines: smooth median (quantile GAM). In (b) the y axis is truncated at 10% for readability, so some extreme values are not shown.

When examined by region, the same temporal boxplots reveal substantial regional disparities in Esca expression levels (Figure 3b). We observe a signal of a negative trend in disease expression with strong interannual variability. The negative trend is quite pronounced in regions with a higher incidence (Chablais, La Côte, Lavaux, Neuchâtel), slight in Valais, and not apparent in northern Vaud where the incidence is quite low. Note that the smooth curves show quite a different behavior, since the median tends to be less sensitive to the interannual variability than the mean. Regional differences were also observed, with esca incidence decreasing more strongly in Chablais, La Côte, Lavaux, and Neuchâtel than in Nord Vaudois and Valais.

### 3.3 FORD algorithm

The FORD *R*^2^ algorithm was applied on both the monthly and seasonal datasets. Among the 42 candidate variables considered in the monthly dataset, FORD identified VPD_July as the sole significant predictor of Esca expression (Figure 4a). In the seasonal dataset, two variables were identified as significant predictors, namely SeasPrecipitation and SeasTemp (Figure 4b). Note that the output of FORD is a set of variables of which the response variable is a function. It does not provide direction of the effect (positive or negative) or a functional form, only a measure of the strength of the association.

**Figure 4.**
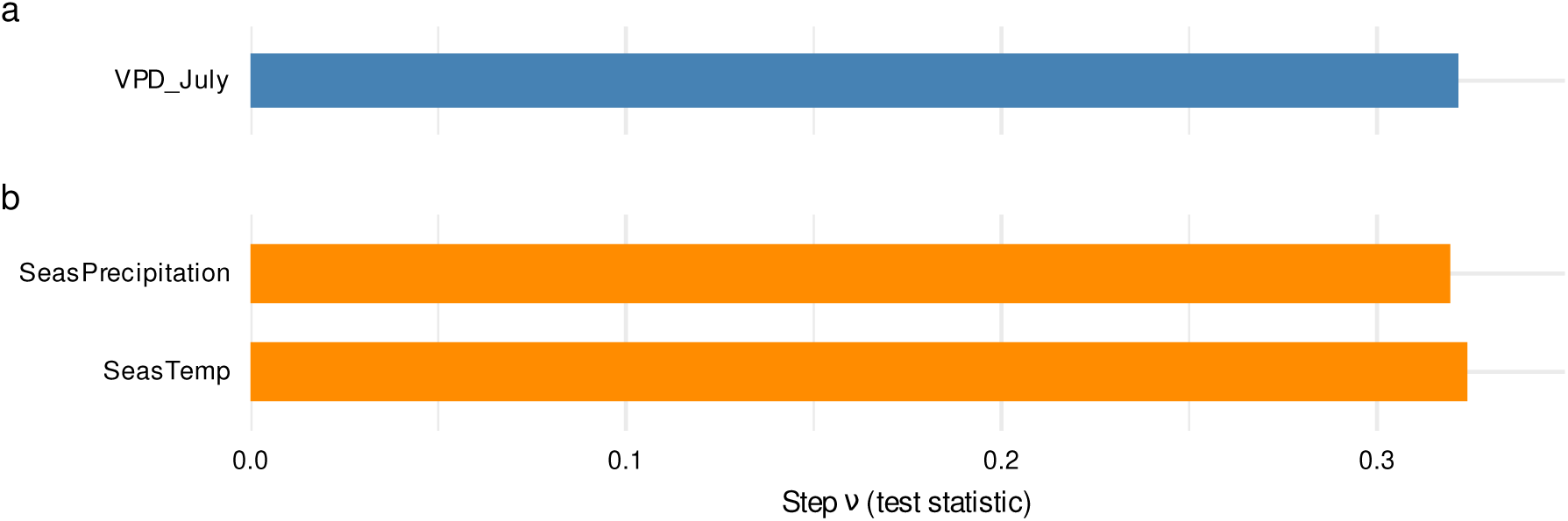
Variables selected by the FORD algorithm, (a) among the monthly climate variables and (b) among the seasonal aggregates. Bars show the test statistic (step *ν*) at each selection step; variables are listed from top to bottom in the order of selection. Only variables with a statistically significant signal beyond random noise are retained: VPD in July for the monthly dataset, and seasonal precipitation followed by seasonal temperature for the seasonal dataset.

### 3.4 Machine learning predictive methods

Across all dataset configurations and models, the predictive performance remained modest. Overall, only the tree-based ensemble methods (random forest, gradient boosting) outperformed the baseline model, but the improvement was moderate, while the LASSO did not beat the baseline (or only slightly).

Figure 5 summarises the model performance in terms of MAE across all models and configurations, including the baseline for reference; the corresponding detailed results are provided in Table S1 in the Supplementary Material.

**Figure 5.**
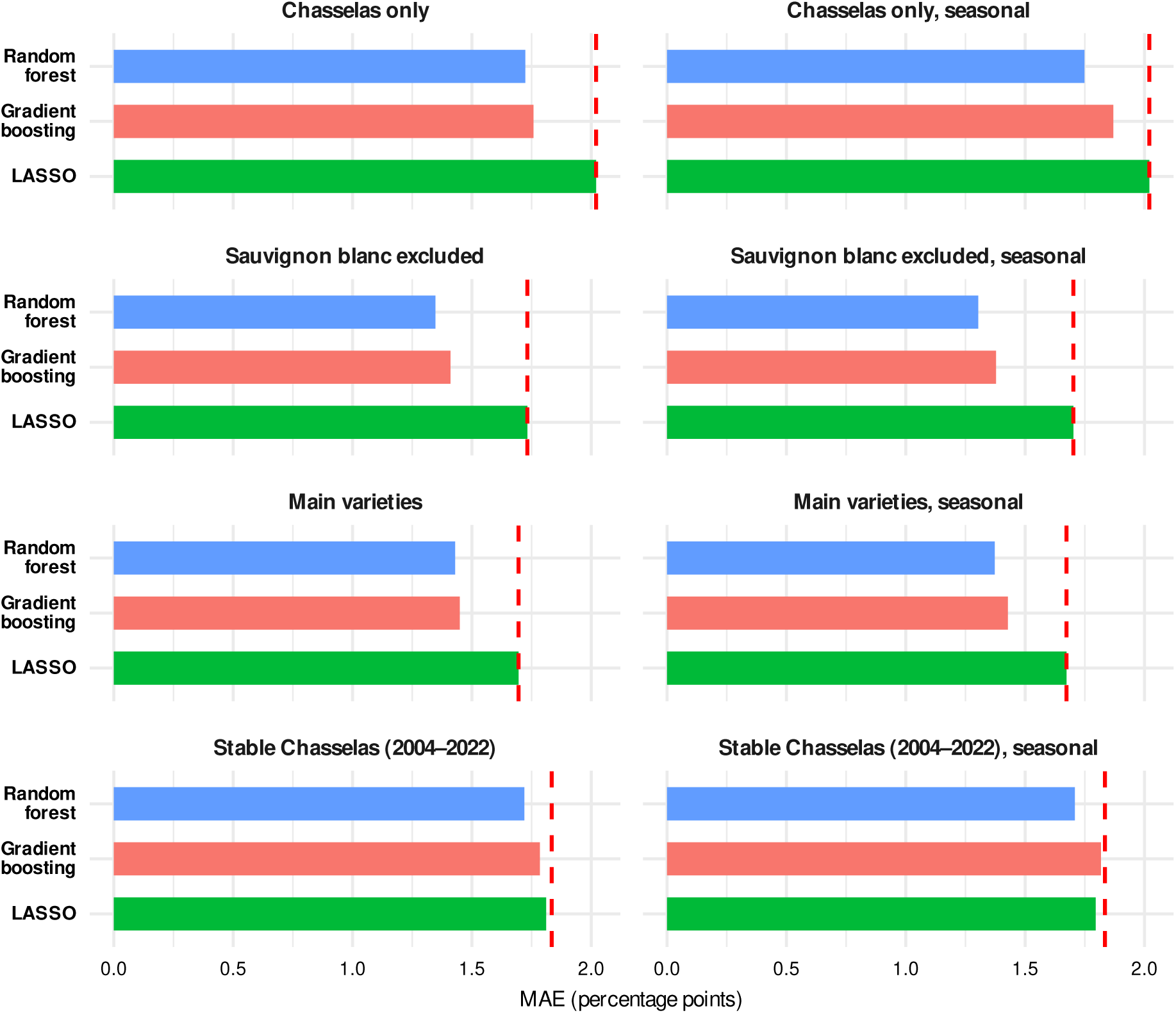
Test-set mean absolute error (MAE, in percentage points) of the three models (random forest, gradient boosting and LASSO) for each data configuration, using monthly climate variables (left) or seasonal aggregates (right). The dashed red vertical line is the baseline MAE (overall mean prediction); bars shorter than the baseline indicate better predictive performance.

We can see that the major gain (23% less of MAE compared to the baseline) is for the random forest trained on the whole dataset but the Sauvignon Blanc. We know that one plot with this variety has a particular history of unexplained susceptibility and is an outlier. The average incidence across all observations is approximately 2.56% (median: 1.5%), so a MAE of ∼1.5 (% percents of incidence) represents a substantial error relative to the average observed incidence.

The best MAE are lower in all the configuration with more than a single cultivar (Exclude Sauvignon and Main varieties). Stable plots or chasselas only do not improve the performance of the models.

We can conclude that using a flexible tree-based model such as random forest and XGBoost improves upon a more interpretable model such as the LASSO. Other, more modern models such as tabPFN (Hollmann et al., 2025) could potentially be used, but we do not expect dramatic improvement.

### 3.5 Variable importance using predictive methods

The combined variable importance analysis, summarized on Figure 6, revealed that the top-ranked variables were reasonably consistent across configurations:

• **VPD** (especially April, May, July, and the seasonal aggregate) was the most consistently top-ranked variable.
• **Atmospheric pressure** (April–July) was prominent across configurations but showed more variability in ranking.
• **Precipitation** (March–May) appeared in the top 10–15 across most configurations.

**Figure 6.**
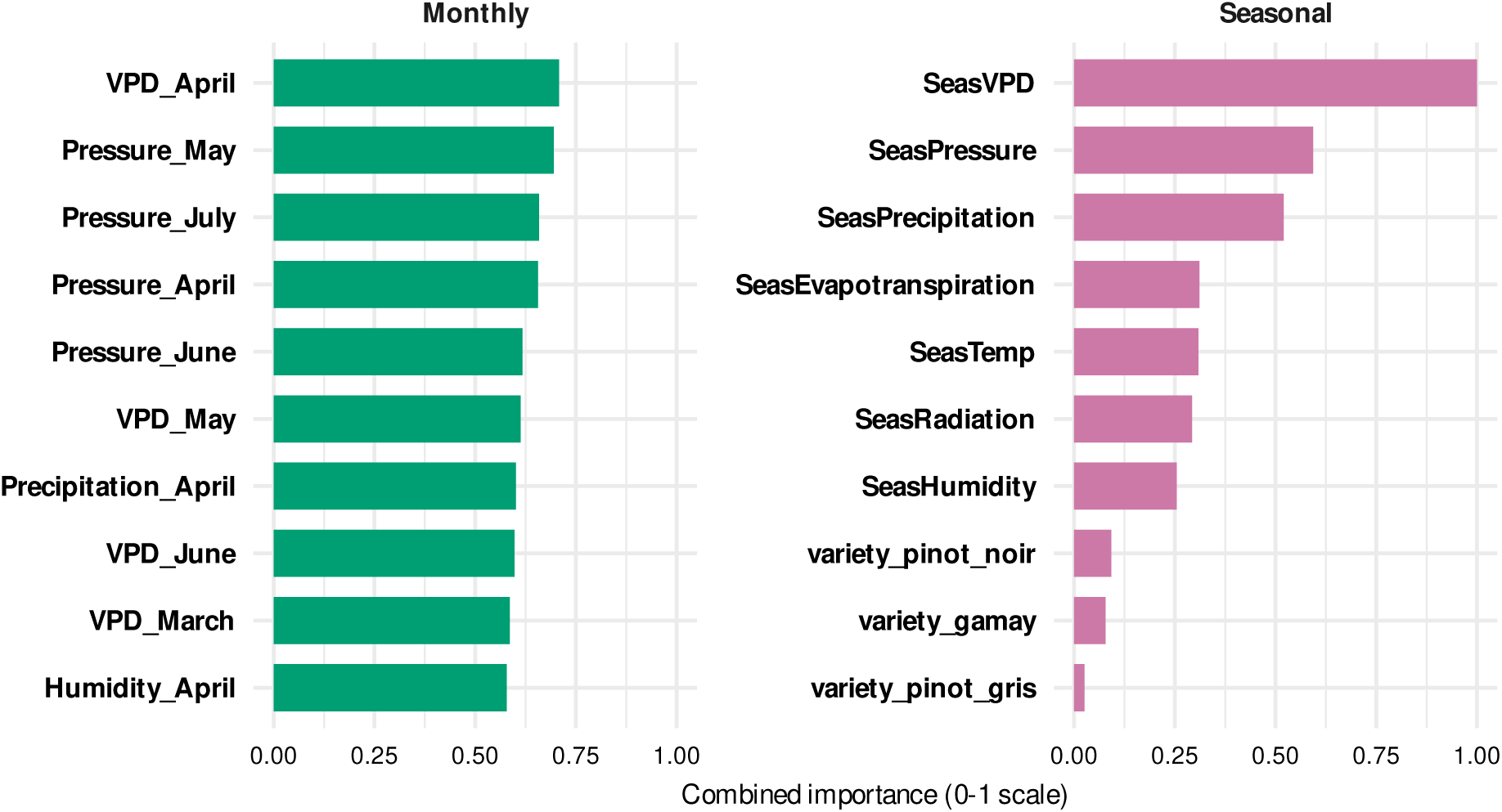
Combined variable importance (0–1 scale) of the ten most important variables, computed across all data configurations and models (random forest, gradient boosting, LASSO) and weighted by model performance, for the monthly (left) and seasonal (right) climate datasets.

The ranking of variables beyond the top 3–5 varied substantially between configurations and models, suggesting limited stability. Relative humidity almost never appears as an important variable. This may be due to its effect being masked by VPD. Radiation also does not seem to affect the incidence of esca disease in any way.

The overall predictive performance remains modest (Figure 5). This limits confidence in fine-grained variable importance distinctions. The results should be interpreted as identifying broad categories of potentially relevant climate drivers (VPD, atmospheric pressure, precipitation) rather than pinpointing specific variables with high confidence.

### 3.6 Regional comparison: rainfall/ET ratio and VPD versus esca incidence

In addition to the predictive study, we sought to examine regional differences through comparisons of hydrometry. Since it has been shown that this factor is critical to the onset of symptoms (Bortolami et al., 2021; Etienne et al., 2026), we also sought to specifically examine this aspect at the regional level. The aim of this aggregation is to reveal a characteristic that is not discernible at plot level and that our predictive analysis fails to highlight. The P/ET ratios varied widely across regions and years, ranging from 0.5 to 2. To better capture the underlying trend, we used a smoothed median (Fasiolo et al., 2021b) which is less sensitive to outliers than a smooth mean.

As shown in Figure 7, higher P/ET ratios, reflecting wetter conditions relative to evaporative demand, and lower seasonal VPD, reflecting lower atmospheric dryness, were associated with higher median Esca incidence.

**Figure 7.**
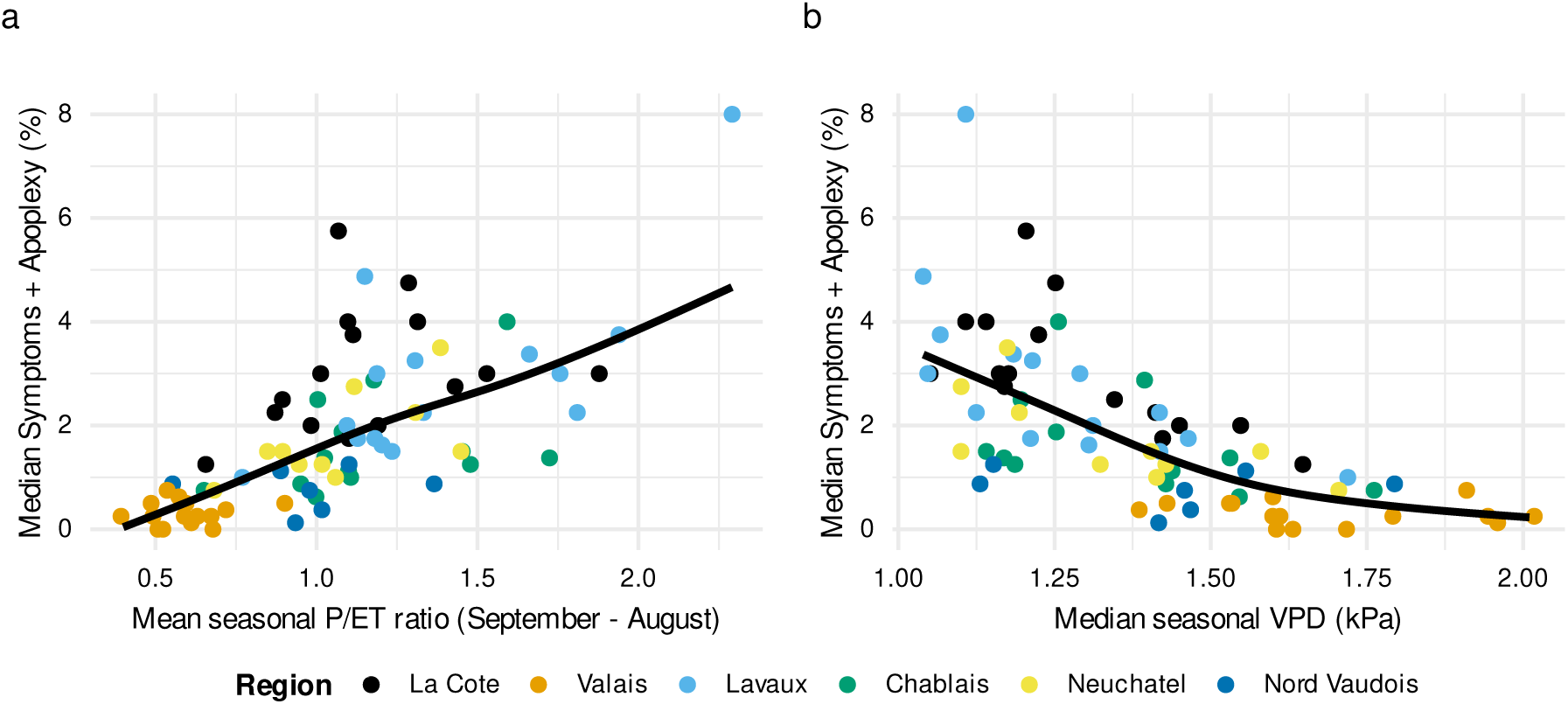
Relationship between esca incidence and climate for each region and year. Each point is a region–year pair; the y axis is the median incidence (% of vine stocks with foliar symptoms or apoplexy) across the plots of the region. (a) Mean seasonal precipitation to evapotranspiration (P/ET) ratio, September–August; (b) median seasonal vapour pressure deficit (VPD, kPa). Colors indicate the region; the black line is a smooth median fitted to all points. Esca incidence increases with the P/ET ratio and decreases with VPD.

Valais, the driest region (P/ET < 1, highest VPD), consistently shows the lowest Esca expression. Conversely, regions such as La Côte and Neuchâtel, which have higher P/ET ratios and lower VPD, show higher Esca expression.

## 4 Discussion

Our 21-year survey revealed substantial spatial and temporal variation in esca incidence across Western Switzerland, with climatic factors related to water balance emerging as important drivers at the regional scale. This network was developed as a surveillance project for regional risk assessment and covers the main viticultural areas of western Switzerland. We analysed the data at two spatial scales: the vineyard level and the regional level.

Over the years of monitoring, esca incidence varied from year to year, both within and across vineyards. This is consistent with the fluctuations in esca incidence reported between years and plots in other long-term monitoring studies (Dewasme et al., 2022; Etienne et al., 2024, 2026). However, we observed a decrease in the overall incidence of esca, contrary to other studies reporting an increase (Bruez et al., 2012) or a stable trend (Etienne et al., 2024). Higher temperature and lower precipitation are considered factors that decrease esca incidence (Bortolami et al., 2021; Etienne et al., 2024) whereas higher precipitation in early summer is positively correlated with esca incidence (Calzarano et al., 2018; Serra et al., 2018). (Scherrer et al., 2022) The increasingly dry conditions observed in Western Switzerland in past two decades, particularly during summer, resulting from changes in precipitation and increased evaporative demand associated with rising temperatures (Scherrer et al., 2022), may have contributed to the observed decrease in esca incidence. These findings highlight the value of long-term monitoring programmes for tracking disease dynamics under changing climatic conditions.

While a decreasing trend was observed across the network, substantial differences emerged among regions. In La Côte, Chablais, Neuchâtel and Lavaux, where esca incidence was reported to be higher, the decreasing trend was more pronounced. Valais and Nord Vaudois, which have historically lower rates of esca incidence, showed a more stable long-term trend. These regional disparities likely reflect differences in climatic conditions and soil characteristics (Calzarano et al., 2018; Marchi et al., 2006; Monod et al., 2025b). Valais is characterized by a drier climate (Zumbrunnen et al., 2009) and well-drained soils (Sigales, 2007) compared to the other regions of our network, which can contribute to explain its lower and more stable esca incidence. In Nord Vaudois, the parent material lies closer to the surface, giving rise to soils with reduced water-retention capacity. Drier climate and poor soil water-retention capacity are therefore factors that reduce esca incidence (Bortolami et al., 2021; Etienne et al., 2026; Monod et al., 2025b). Interestingly, substantial variation was also observed among plots within the same wine-growing region, despite comparable climatic conditions. This suggests that, in addition to the high inherent randomness, local factors such as pedological features may play a critical role in disease expression.

We assessed the predictive power of several models using meteorological variables to predict esca incidence (% of symptomatic plants per plot) across our plot network, but the performance remained poor at the plot level. The low prevalence of symptomatic vines (approximately 2-3%) combined with relatively limited interannual variation reduced the signal available to modelling approaches, making accurate predictions inherently challenging. Esca incidence cannot be explained by meteorological variables alone, in line with previous studies conducted in various geographical contexts (Bruez et al., 2012; Dewasme et al., 2022; Etienne et al., 2026; Lecomte et al., 2024). These findings suggest that intrinsic plot characteristics may contribute more strongly to disease expression than meteorological variables alone, suggesting that other, still unidentified factors are also involved.

The output of the FORD algorithm shows that, at the monthly scale, VPD was the variable most consistently associated with esca incidence, whereas at the seasonal scale, precipitation and temperature showed the strongest associations. Among all variables tested, VPD also appeared to be the most stable predictor across models, which is known to be an important driver of plant functioning (Grossiord et al., 2020). Although the overall predictive performance of the models remained modest, these findings are consistent with previous studies. For example, Etienne et al. (2026) reported a positive relationship between temperature and precipitation during the symptom expression period and a negative relationship between VPD during the growing season and esca incidence. Similarly, VPD, temperature, and precipitation have been identified as relevant climatic drivers of esca expression in other viticultural contexts, lending further support to our results. These variables are closely linked to seasonal dryness, which is increasingly recognized as a major determinant of esca incidence (Bortolami et al., 2021).

Under drier conditions, elevated VPD increases atmospheric water demand and promotes stomatal closure, thereby reducing transpirational water loss (Grossiord et al., 2020) and hydraulic conductance along the soil to leaf pathway (Zufferey and Smart, 2012). One possible explanation is that reduced xylem transport under water-limited conditions could limit the movement of phytotoxic compounds involved in symptom expression, although experimental evidence supporting this mechanism remains limited (Bortolami et al., 2021). Conversely, greater water availability may facilitate transpiration and xylem transport processes, potentially increasing the movement of such compounds within the plant.

Since our models did not achieve strong predictive power for esca incidence based on meteorological variables at the plot level, we performed regressions using the most promising variables (VPD and the precipitation-to-evapotranspiration ratio) against esca incidence by region. At the regional scale, a higher precipitation-to-evapotranspiration ratio was associated with an increase in esca incidence, which in turn are generally associated with greater vine vigor. In addition, higher precipitation is likely associated with increased soil water availability, reflected by a higher Soil Water Index (SWI) which is a determinant factor known to increase esca incidence (Etienne et al., 2026; Monod et al., 2025b). At the other end of the gradient, Valais, the driest region considered, consistently showed the lowest esca incidence. The correlation between high VPD and lower esca incidence across regions confirms this pattern: when atmospheric demand is higher, meaning a greater water deficit, esca incidence is lower. Since the regions covered by our dataset exhibit a wide range of hydrological regimes, our data set provide a reasonably uniform coverage of the interval of both ET/P and VPD variables. This regional-scale pattern is consistent with the variable importance and FORD results: VPD and water-related variables are the dominant climatic drivers of Esca expression. The relative concordance between the plot-level statistical analyses and this simple regional characterization reinforces the conclusion that water stress (or lack thereof) is the primary climatic factor modulating Esca symptom expression that our data can identify. This strengthens our confidence in the association between these variables and esca incidence. The growing body of evidence linking water availability with esca incidence suggests that irrigation practices should be carefully considered, particularly in susceptible cultivars. (Etienne et al., 2026).

One of the key findings of this study is that meteorological variables explained variation in esca incidence more effectively at the regional than at the plot scale. This suggests that climate may primarily influence esca incidence at a broader spatial scale while local environmental, pedological, and biological factors may play a greater role in determining disease expression at the vineyard level.

## Acknowledgements

These data represent hundreds of hours of fieldwork and thousands of kilometers travelled over more than two decades. We are deeply grateful to all those who contributed to data collection over the years, in particular Nicole Lecoultre, Sylvain Schnée, Éric Remolif, Bernard Bloesch, Isabelle de Groote, Jean Taillens, and Aurélie Melgar.

## Supplementary material

The whole analyses are available on a public GitHub repository and are reproducible. The data set has been made available on Zenodo.

**Table S1.**
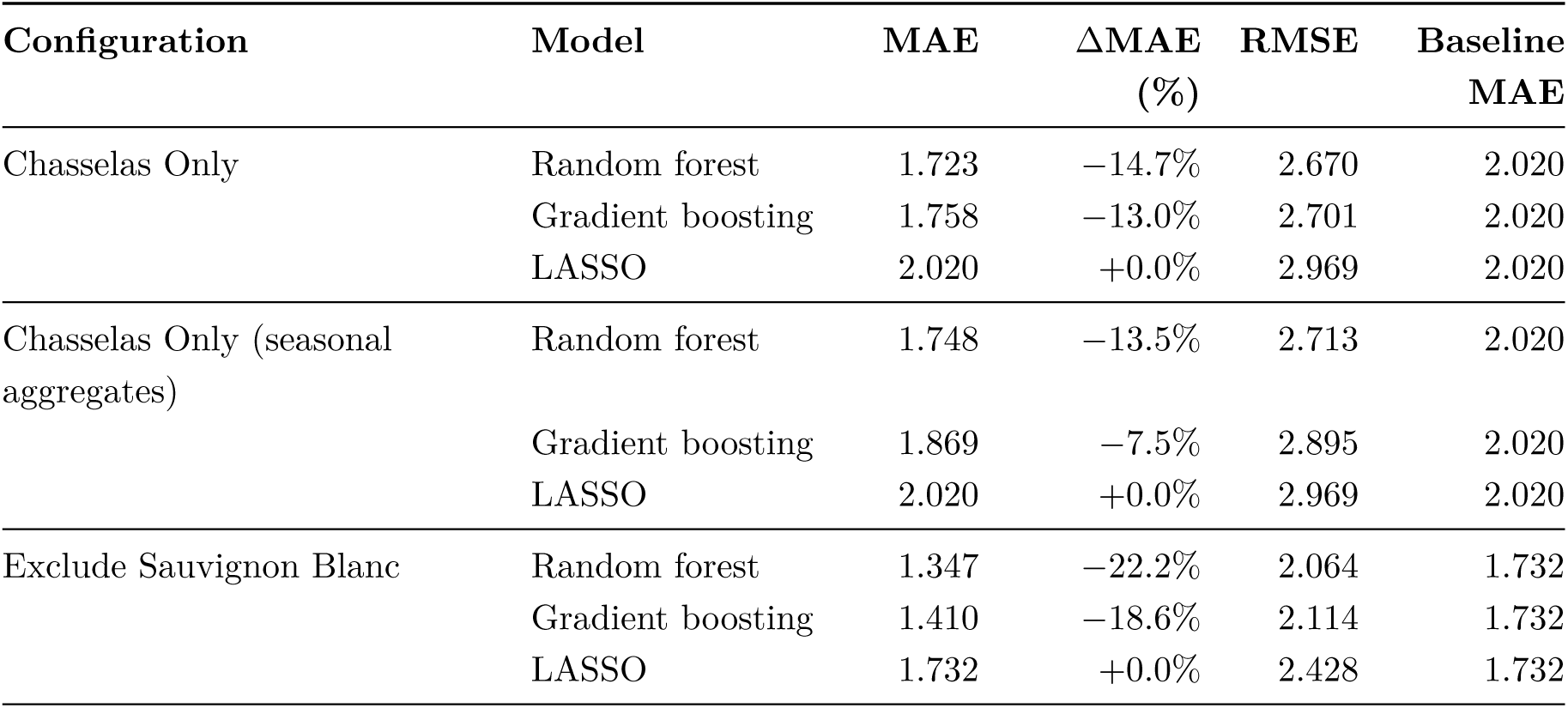
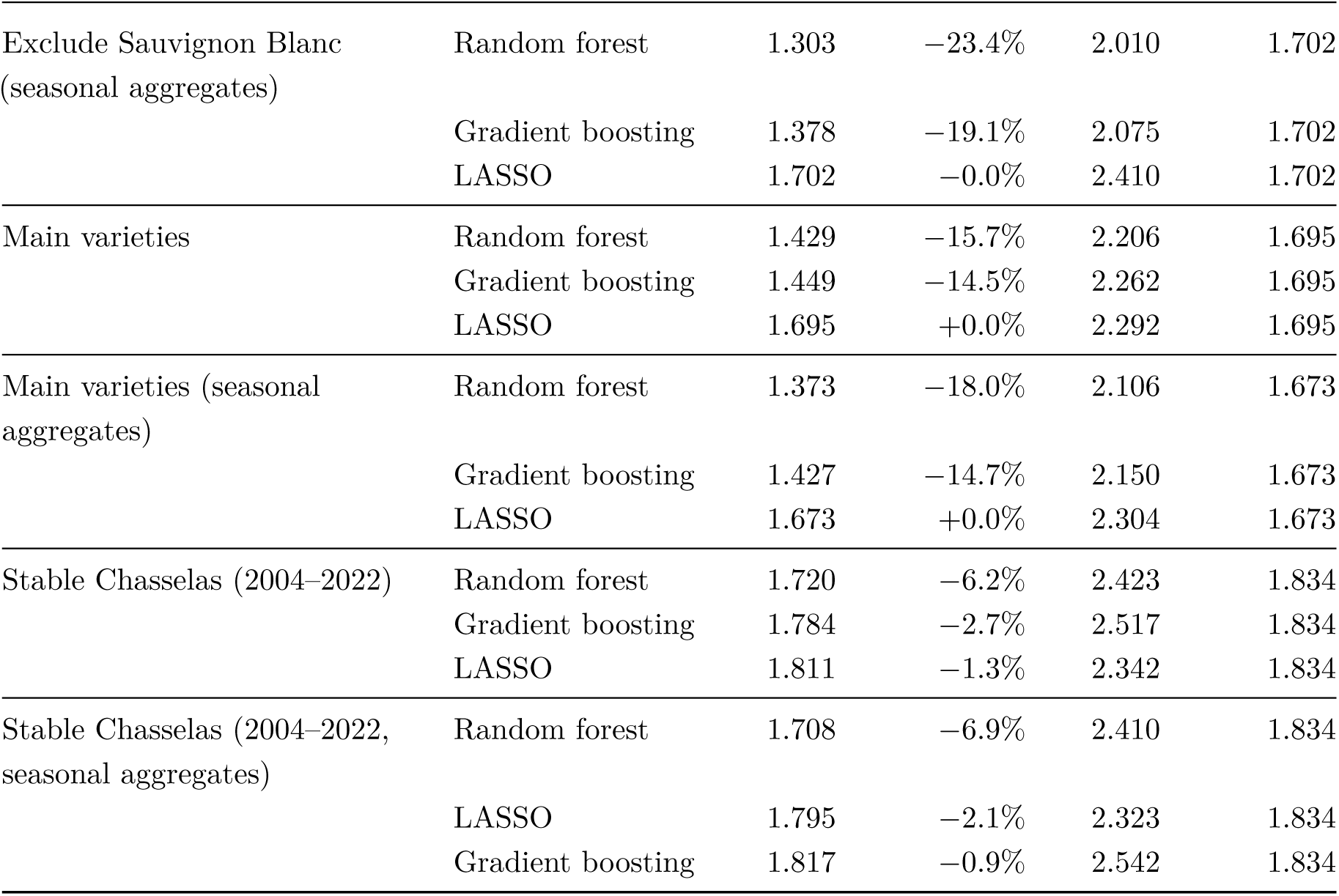
Performance of the predictive models (test set). MAE: mean absolute error; RMSE: root mean squared error; both in percentage points. ΔMAE is the relative difference of the MAE with respect to the baseline MAE.

